# Learning a conserved mechanism for early neuroectoderm morphogenesis

**DOI:** 10.1101/2023.12.22.573058

**Authors:** Matthew Lefebvre, Jonathan Colen, Nikolas Claussen, Fridtjof Brauns, Marion Raich, Noah Mitchell, Michel Fruchart, Vincenzo Vitelli, Sebastian J Streichan

**Author notes:** These authors contributed equally (Matthew Lefebvre, Jonathan Colen, Nikolas Claussen).

## Abstract

Morphogenesis is the process whereby the body of an organism develops its target shape. The morphogen BMP is known to play a conserved role across bilaterian organisms in determining the dorsoventral (DV) axis. Yet, how BMP governs the spatio-temporal dynamics of cytoskeletal proteins driving morphogenetic flow remains an open question. Here, we use machine learning to mine a morphodynamic atlas of Drosophila development, and construct a mathematical model capable of predicting the coupled dynamics of myosin, E-cadherin, and morphogenetic flow. Mutant analysis shows that BMP sets the initial condition of this dynamical system according to the following signaling cascade: BMP establishes DV pair-rule-gene patterns that set-up an E-cadherin gradient which in turn creates a myosin gradient in the opposite direction through mechanochemical feedbacks. Using neural tube organoids, we argue that BMP, and the signaling cascade it triggers, prime the conserved dynamics of neuroectoderm morphogenesis from fly to humans.

## INTRODUCTION

Morphogenesis is the process by which the shape of an organism emerges from the coordinated behavior of groups of cells. Turing’s pioneering work traced morphogenesis to the presence of “chemical substances called morphogens, reacting together and diffusing through a tissue” [1]. Molecular biology has since identified these morphogens as molecules whose concentration modulates the expressions of various genes [2]. During early development, morphogen concentrations set up a spatial coordinate system and body axes on which the embryo organizes future tissues and organs [3]. However, it is force-generating proteins – not morphogens – that produce the active mechanics which moves each group of cells into the right place [4–6]. This raises the questions: *How do morphogens control the active mechanics that execute morphogenesis? And are these mechanisms conserved across species?*

We focus on the paradigmatic example of neuroectoderm morphogenesis, which is the first step in forming the central nervous system in species ranging from *Drosophila melanogaster* to *Homo sapiens*. This process is regulated by bone morphogenetic protein (BMP) signaling, whose role as a morphogen is conserved across organisms with bilateral symmetry [7, 8]. In addition, tissue elongation is driven by a second conserved mechanism known as convergent extension [9]. However, the exact process by which BMPs control tissue mechanics is unknown.

The dynamics of these tissues is dauntingly complex. Nonetheless, a remarkable simplification occurs when focusing on large scale patterns. To obtain consistent and reproducible dynamics across different embryos, we coarse-grain data, i.e. we smooth it in space and align it in time [10]. Recent advances in light-sheet microscopy [11] have enabled the collection of such coarse-grained data for tissue and protein dynamics across the entire embryo [10]. At the cellular level, tissue deformations arise from the interaction of many biomolecules, as exemplified in Box 1 for *Drosophila*. At the coarse-grained level, a smaller number of these correlated quantities may be sufficient to predict the large-scale mechanical behavior. In order to identify the minimal biochemical information that allows one to encode tissue flow, we use statistical inference tools such as machine-learning (ML) [12–14], as shown in Fig.1e. These techniques alone are not sufficient to learn causal rules for tissue dynamics. To do so, we must supplement ML with insights from biology and physics [15–17]. Here, we develop a pipeline that integrates *in toto* light-sheet microscopy, interpretable machine learning, and physical theories of active mechanics in order to develop and test a biological hypothesis: BMP shapes the dynamics of convergent extension by initiating mechanochemical feedback loops involving force-generating proteins as well as genetic regulation.

This paper is organized as follows. We first analyze convergent extension in *Drosophila* embryos. Using our integrated pipeline, we identify predictive biophysical quantities — tissue flow, myosin, and adherens junctional proteins — and causal rules which describe their coupled dynamics during a key tissue deformation that occurs during Drosophila development, known as germ-band extension (GBE) [18]. We then test the validity and generalizability of these rules using mutant analysis. In a nutshell, BMP establishes a global cell adhesion pattern which modulates mechanochemical feedbacks to produce a myosin contractility gradient in the opposite direction. Finally, we turn to experiments on human stem cell-based neural tube organoids and unveil how BMP, and the signaling cascade it triggers, prime the conserved dynamics of neuroectoderm morphogenesis from flies to humans.

### Box 1.

***Drospohila* morphogenesis: a primer**

In *Drosophila*, convergent extension drives a major morphogenetic movement known as germ-band extension (GBE) [18], in which the neuroectoderm elongates along the anterior-posterior (AP) axis (*right*, a-c). This process arises through organized cell intercalations, in which cell interfaces oriented along the dorsal-ventral (DV) axis contract and are replaced by ones oriented along the anterior-posterior (AP) axis (*right*, d). The mechanics at cell interfaces (*right*, b) is driven by the interplay between force-generating proteins such as non-muscle myosin-II (myosin), a motor protein which causes junctions to contract, and the adherens junction complex which is involved in building the substrate for myosin. It has been proposed that transcription factors known as pair-rule-genes (PRGs) organize germ-band extension by regulating a family of receptors, known as Toll-like receptors [19, 20], located at cell junctions (*right*, e). PRGs are patterned in periodic stripes along the AP axis (*right*, e), and could therefore provide a directionality to the process by modulating the recruitment of force-generating proteins depending on the orientation of their interfaces. On the other hand, it has been shown that junctional myosin contraction exhibits a gradient along the DV axis, which is sufficient to predict the tissue flow during convergent extension [21]. However, this global myosin gradient is stationary, and cannot be attributed to PRG patterns that rotate with the tissue [22] but can be explained by mechanochemical feedback loops [23, 24]. The outstanding challenge therefore becomes: how are the AP and DV axes integrated? Understanding the dynamics of myosin holds the key to address this question.

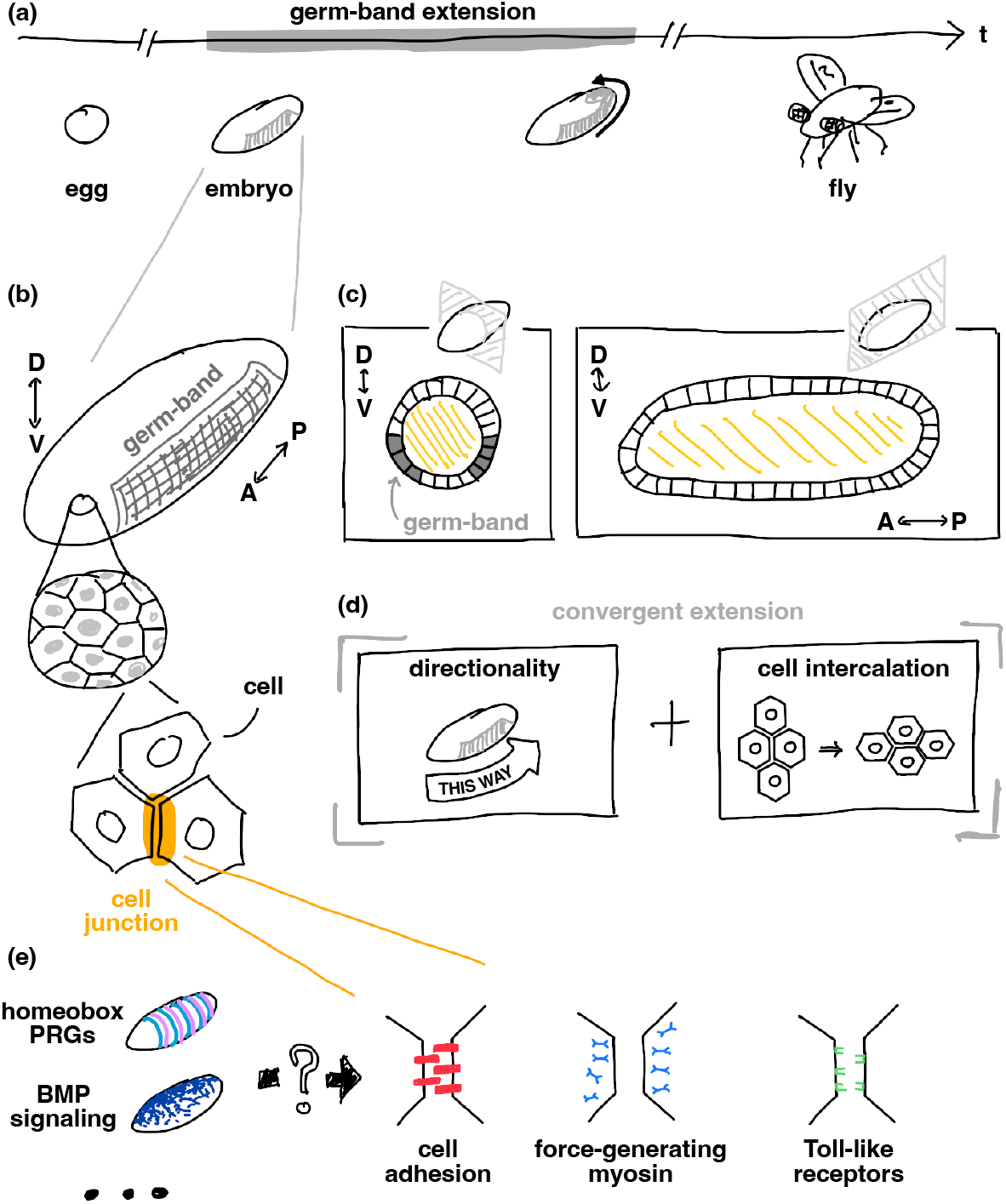

## RESULTS

### Neural networks forecast tissue dynamics from initial myosin distribution

We first aim to isolate the smallest set of proteins that allow us to forecast mechanical behavior from initial conditions. To do so, we leverage a morphodynamic atlas of *Drosophila* gastrulation which contains *in toto* protein expression movies for various patterning genes and components of the cytoskeleton [10] (Fig. 1a-e). We then train deep neural networks (NNs) to learn tissue dynamics (see SI for details) using different combinations of these movies. This agnostic approach allows us to identify predictive biochemical quantities without specifying a physical or biological model beforehand (Fig. 1e-f). For our purpose, it is actually advantageous that neural networks are essentially input-output black boxes agnostic to the biological rules that we are yet to discover. We find that knowledge of the initial myosin distribution (*t* = 0 on Fig. 1a corresponding to ventral furrow formation) is sufficient for a NN to learn to forecast tissue flow in wild type (WT) embryos for 20 minutes and maintain excellent agreement with experiments through the onset of GBE (Fig. 1g). Previous work has evidenced an instantaneous relation between myosin and tissue flow during convergent extension [21]. Our results go further, suggesting that myosin contains enough information to quantitatively account for the dynamics of cellular flow and resulting trajectories (Fig. 1g). Finally, comparison of NNs trained on different sets of proteins indicates that the myosin distribution is necessary to forecast the flow from an initial condition: Fig. 1h shows that any other protein in the atlas (or combination thereof) produces a higher prediction error.

**Figure 1:**
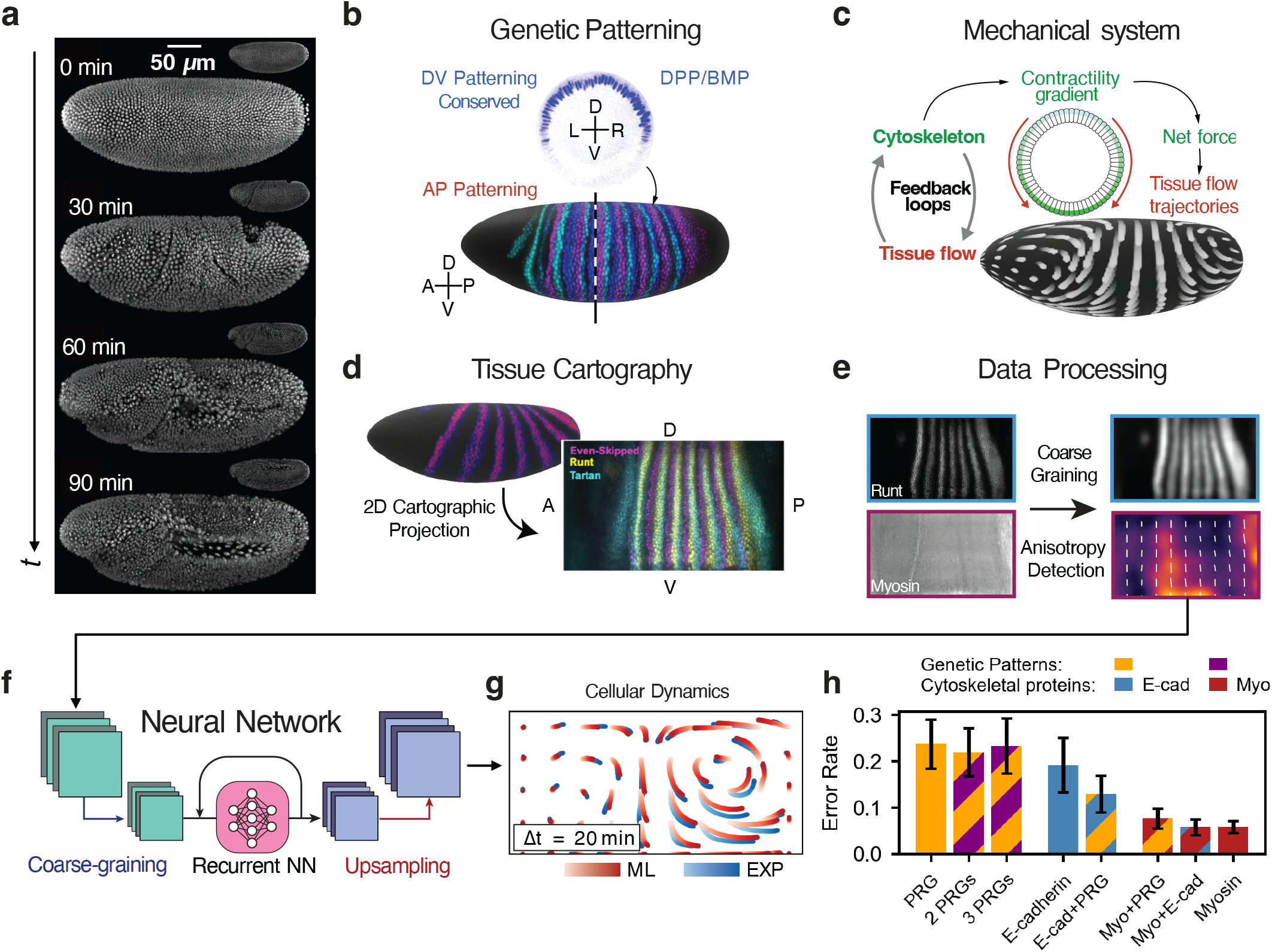
Drosophila GBE as a system for understanding the interplay between genetic patterning and cytoskeletal force generation using machine learning. (**a**) Snapshots of nuclei from *in toto* live recording of 90 minutes of *Drosophila* gastrulation, comprising invagination of the ventral furrow and GBE. (**b**) The conserved morphogen DPP patterns cell fates along the DV axis, fates illustrated in blue. The AP axis is patterned by striped expression of PRGs, three of which are shown (Even-skipped: blue, Runt: cyan, Paired: magenta, virtual overlay created from 3 different embryos). (**c**) A gradient of myosin recruitment along the DV axis generates forces that drive tissue flow. Mechanical feedback loops respond to tissue deformation, sculpting local patterns of cytoskeletal recruitment. (**d**) Tissue cartography projects the curved 3D embryo surfaces onto a 2D plane. (**e**) Additional image processing includes spatial downsampling/smoothing and optional anisotropy detection via a radon transform algorithm. Anisotropy detection yields a tensor field, which can be thought of as a double-headed arrow whose direction aligns with cell junctions where a protein lies and whose magnitude reflects the local protein density. (**f**) A NN forecasts tissue dynamics from the measured biological initial conditions. The network is a residual autoencoder which maps the inputs to a latent vector, predicts dynamics using a recurrent layer, and then translates the latent vector sequence into flow fields. (**g**) Trajectories of test points in flow fields measured in experiment (blue) and predicted by NN from an initial snapshot of myosin (red). (**h**) Forecasting performance of NNs trained on genetic and cytoskeletal patterns. NNs trained on myosin achieve the lowest error rates.

### Coarse-grained dynamics of GBE are low-dimensional and reproducible across embryos

Neural networks can perform forecasting, but are ultimately a black box from which it is difficult to infer interpretable causal rules. To gain further insight into the dynamics of myosin and flow during GBE, we perform a principal component analysis (PCA) of the coarse-grained fields and track the time evolution of the weight of each principal component over the entire duration of GBE (Fig. 2a). In a nutshell, PCA decomposes the data into characteristic features known as “principal components” (PCs). The weight of a PC describes how much this component contributes to the data. As illustrated in Fig. 2b, we consider a fixed set of PCs and encode the time evolution of the data into time-dependent weights *a*_*i*_(*t*) of the PCs *PCi*. We find that a small number of PCs can account for the dynamics of proteins and tissue flow during GBE (Fig. 2c). Moreover, the dynamics in the low-dimensional spaces represented by the primary PCs – the dominant patterns – are simple and repeatable across embryos (Fig. 2a, fourth column). The PCs give us insight into the dynamics: there is little myosin at early times, but the weight of the first PC increases as gastrulation proceeds, indicating development of a DV-patterned myosin field. Similarly, there is little flow at early times, but the weight of the first PC increases during the onset of GBE where a characteristic flow pattern with four vortices develops. The simplicity and reproducibility across embryos revealed by PCA suggest that simple rules for GBE can be obtained.

**Figure 2:**
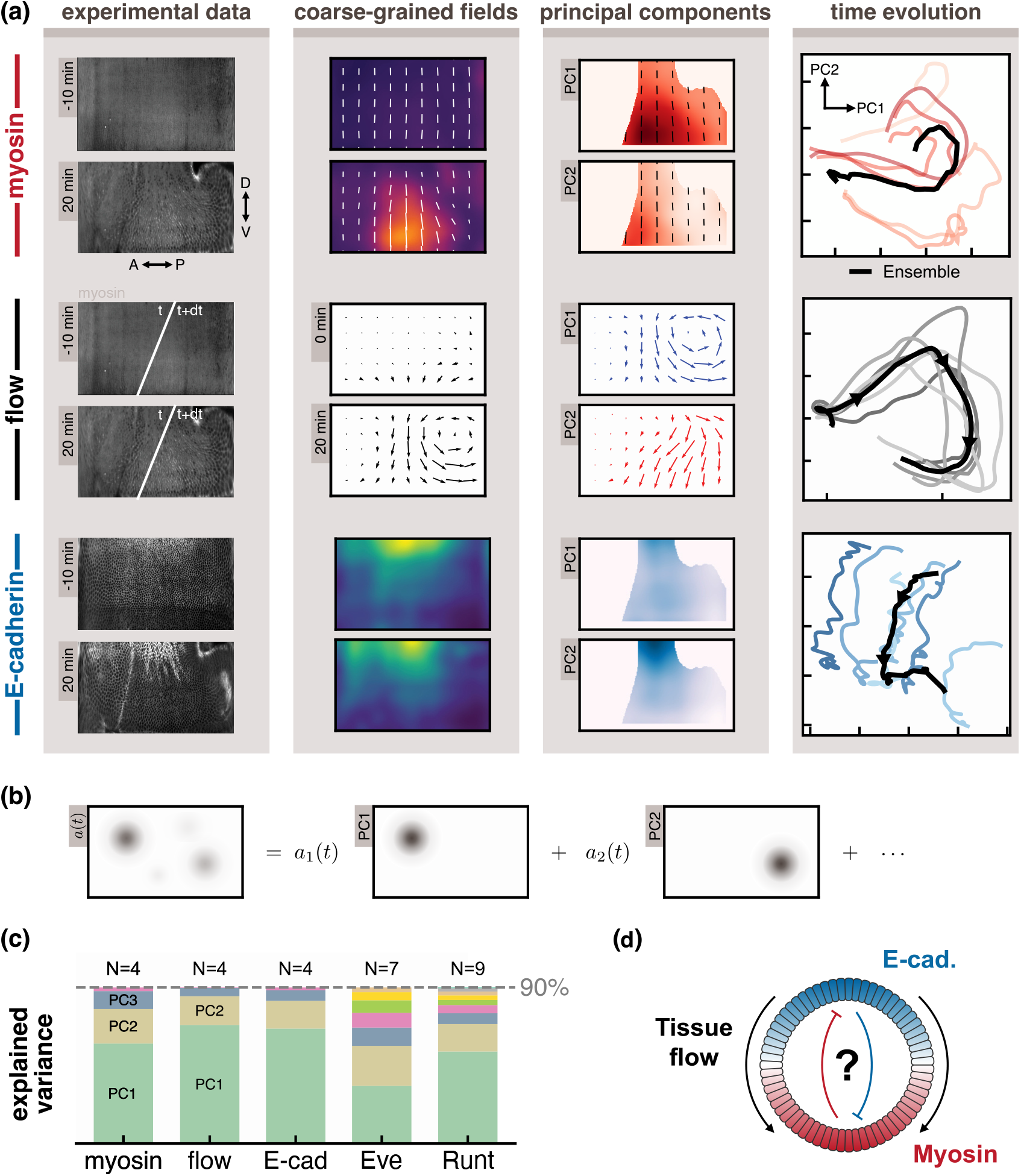
Coarse-grained dynamics of GBE are low-dimensional and reproducible across embryos. (**a**) (first column) Representative fluorescence images from experimental movies. Times are relative to initiation of the ventral furrow at *t* = 0 min. Flow is computed via PIV on the fluorescence images. (second column) Myosin is coarse-grained using a radon transform algorithm into a tensor field that captures density and junctional orientation. Tissue flow is a vector field obtained via PIV. E-cadherin is coarse-grained into a scalar density field by smoothing the fluorescence image. (third column) Primary PCs for myosin and E-cadherin are DV-graded in opposite directions. Here we mask out cephalic furrow and posterior midgut regions which become prominent at later times. Primary PCs for tissue flow describe the positions of vortices in the GBE flow field. (fourth column) Projection of dynamics onto the space spanned by the primary PCs demonstrates smooth and reproducible behavior across embryos. (**b**) PCA decomposes a complex pattern into a number of characteristic features, or principal components. (**c**) We quantify the complexity of a field by the number of principal components required to describe 90% of dataset variation. Myosin, tissue flow, and E-cadherin require 4 PCs, while PRGs (Runt and Even-Skipped) require more. (**d**) The relation between the PCA of myosin and E-cadherin suggests their dynamics may be coupled.

### Active matter models of GBE require a patterned control field

Having identified the most crucial features of the myosin pattern and tissue flow, we attempt to reproduce their dynamics using minimal mathematical models inspired by active matter theories that successfully describe cell and tissue mechanics [4, 5, 25].

It is challenging to find a dynamical system including only myosin and flow that reproduces long-lived dynamic states of development observed in experiments (Fig. 2a).The challenge mainly consists in obtaining a myosin gradient from initially nearly uniform myosin and velocity fields (at *t* = 0, see Fig. 2a). As myosin is always higher on the ventral pole, we conclude that it must be modulated by some field that exhibits a gradient at early times (and exclude a pattern-forming instability). Such a “control” field, which may be established by upstream regulation, induces a myosin gradient along the DV axis, which in turn causes a GBE-like flow with four vortices. In addition, we find that a mechanical feedback loop that couples tissue deformation to myosin recruitment (see Box 1 and Refs. [21–23, 26, 27]) can extend the lifetime of this quasi-stationary state, which would otherwise be destroyed by nonlinear effects such as advection and junction rotation.

### E-cadherin acts as a patterned control field for GBE

We now aim to identify a biochemical candidate for the control field that establishes the myosin recruitment gradient. As shown in figure 2a, E-cadherin expression harbors some of the features expected of such a control field. In particular, E-cadherin exhibits a DV gradient before the onset of GBE (Fig. 2a, bottom row), and can therefore serve as a control field for GBE. The mechanics of tissue flow arises from the behavior of cell junctions, where myosin is recruited and generates forces (Box 1). E-cadherin couples adjacent cells to each other at cell junctions, and is critical to establish the substrate to which the actin cytoskeleton is anchored. Motivated by the observation that other junctional proteins have similar expression profiles, we propose that E-cadherin can be used as a proxy to represent the overall state of cell junctions. In addition, high levels of E-cadherin have been correlated with an inhibition of junctional recruitment of myosin [28].

At the same time, we find that the number of PCs needed to represent the dynamics of E-cadherin is identical to that of myosin and tissue flow (Fig. 2c). The relation between the PCA of myosin and E-cadherin suggests their dynamics may be coupled (Fig. 2d). The primary PCs of E-cadherin are DV-graded in the opposite direction of myosin (Fig. 2a) and undergo similar dynamics. We note that in other contexts, morphogen gradients in opposite directions are known to shape stable gene expression patterns [29–35].

We could also have considered other upstream regulators such as PRGs (Box 1). Indeed, mutations to certain PRGs lead to abnormal myosin recruitment and tissue flow during GBE [18, 19, 22, 26]. However, we find that PRG patterns contain much more complex information (Fig. 2c). This can be traced to the observation that PRGs flow with the tissue: their stripes continuously change their orientation, leading to an orientation discrepancy with myosin, whose orientation is nearly stationary [22]. This motivates us to first focus on a post-translational picture.

### Machine learning yields interpretable rules for neuroectoderm morphogenesis

At this stage, our tentative picture is as follows: E-cadherin acts as a control field to recruit myosin, which puts cells into motion. We now aim to assemble these ingredients into a mathematical model that describes experimental data. Using a ML technique known as SINDy [36], we learn interpretable biophysical equations for the coupled dynamics of myosin and E-cadherin (see Box 2). These equations represent rules and feedback mechanisms which ultimately govern *Drosophila* neuroectoderm morphogenesis. These rules involve several mechanisms that we traced back to existing experiments: myosin detachment [22], tension recruitment [22, 24], and strain-rate feedback [23]. Crucially, we find that the strength of each of these mechanisms is modulated by the local E-cadherin concentration.

To evaluate these dynamical equations, we must also model the instantaneous relation between myosin and flow [21]. Here, we use a neural network in order to account for external forces due to the ventral furrow invagination, another part of *Drosophila* morphogenesis that we do not analyze (Fig. 3a-b). Our hybrid model predicts the development of a dorso-ventral myosin gradient opposite to the E-cadherin gradient (Fig. 3c-e) and the transition to a vortical flow pattern (Fig. 3c). We accurately forecast the developmental trajectory as a closed loop for up to 30 minutes, starting 10 minutes before initiation of the ventral furrow, and ending during GBE (Fig. 3b). How does the anisotropy of the myosin nematic field emerge? Based on our model, this anisotropy originates from two sources: first, the geometry of the cell explicitly breaks the symmetry (this is modeled by the geometric stress term in Box 2); second, the formation of the ventral furrow pulls on the embryo (leading to the active tension and strain-rate recruitement in Box 2).

**Figure 3:**
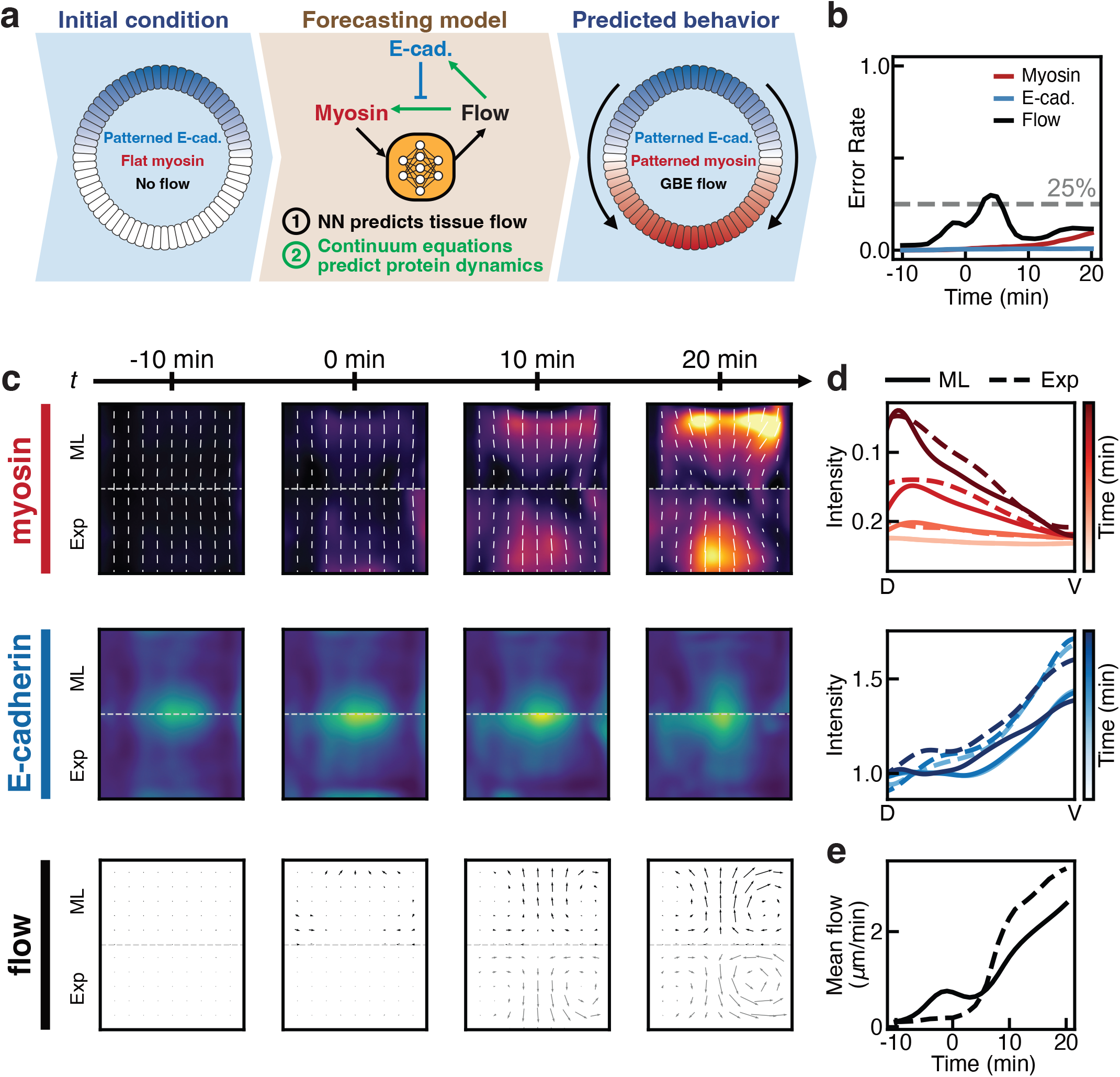
Predicting dynamics of cytoskeletal proteins and flow. (**a**) Schematic of machine-learned dynamical system. At each time step, a NN (Fig. 1f) predicts instantaneous tissue flow from myosin. Eqs. 1-2 (see Box 2) predict the time derivatives of the myosin and E-cadherin fields, which are integrated to forecast embryo behavior from initial conditions. (**b**) Error rate of predictions over 30 minutes, beginning 10 minutes before VF. The error for each field remains within ≈25% over the trajectory. (**c**) Snapshots of predicted myosin, E-cadherin, and tissue flow at 10-minute increments. The top half of each row shows the model prediction (ML) while the bottom half shows the coarse-grained fields measured in experiment (Exp). (**d**) DV variation in model predictions (solid line) and experiments (dashed line) over time for myosin and E-cadherin. Myosin develops a DV gradient from uniforms initial conditions, while E-cadherin maintains an initial DV gradient. (**e**) Average flow magnitude predicted by the model (solid line) and measured in experiment (dashed line). The model predicts the onset of GBE flow arising from the developing DV contractility gradient.

#### Box 2.

**Learning an interpretable model of embryo development**

After considering minimal models, we use the SINDy method to learn rules from the data itself. In brief, this method constructs dynamic equations from a large *library* of possible terms, with the aim of fitting the data with as few terms as possible (see SI for details). Eq. 1 is a passive advection equation for E-cadherin. The cartoons below Eq. 2 are visual interpretations of each term. The LHS is constrained to the co-rotational derivative for the given velocity field. The first RHS term captures the propensity for myosin motors to detach from junctions. The second is tension recruitment due to local myosin-driven active stress and the third is an embryo-scale static stress. Finally, there is a mechanical feedback which recruits myosin to strained junctions.

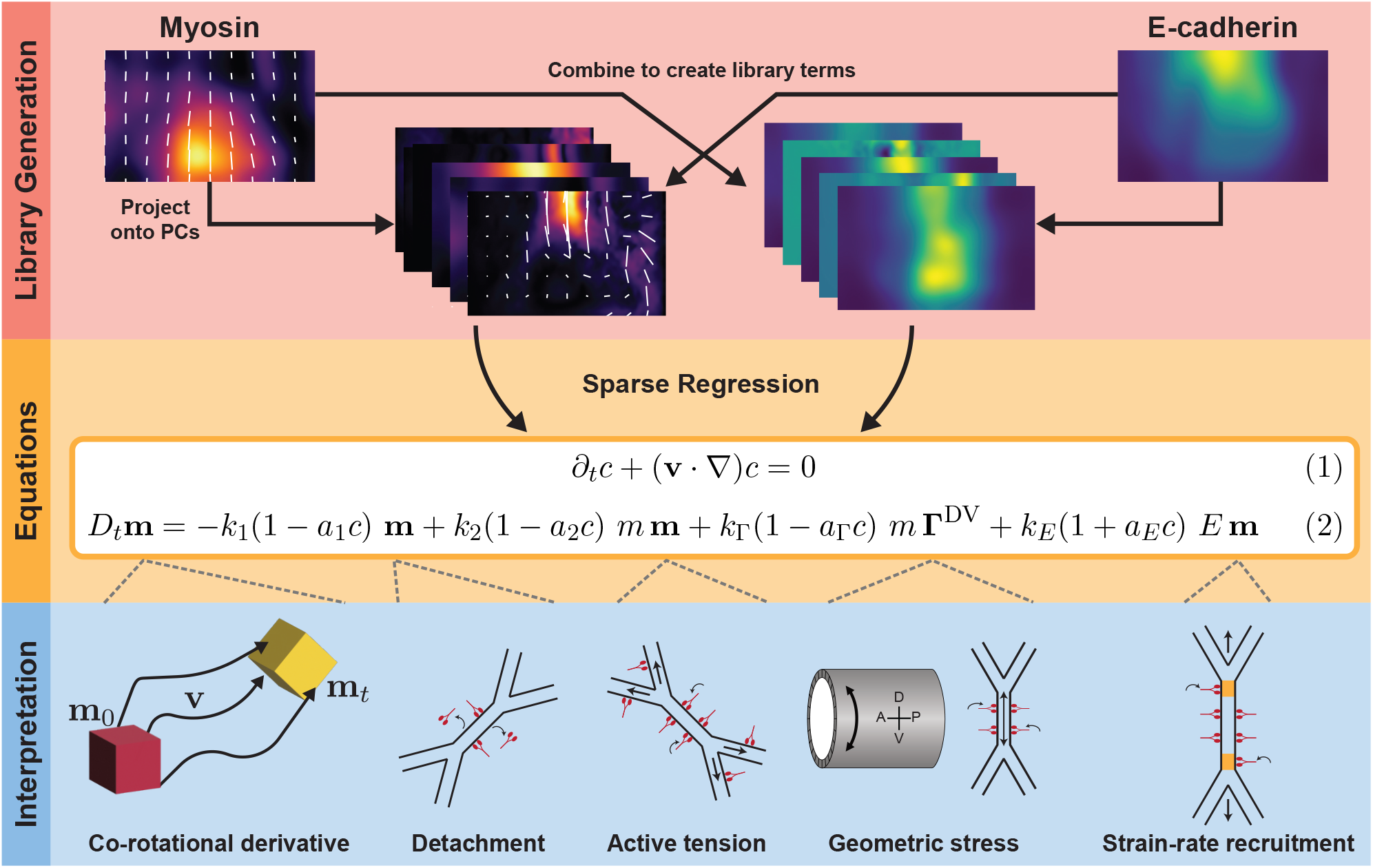

Each mechanism is modulated by the cadherin pattern *c*. The (1 − *c*) prefactors establish a myosin DV-gradient opposite cadherin. The strain-rate recruitment term increases myosin concentration as a tissue is stretched. We note that the coarse-grained field **E** cannot distinguish contributions from two distinct biological processes, cell rearrangement and cell stretching [37]. Biologically, only the latter should recruit myosin. The learned strain rate term has a (1 + *c*) prefactor, which we interpret as distinguishing the strains from these two sources. Rearrangement dominates laterally while deformation primarily occurs dorsally, correlating with the cadherin pattern. Thus, the macroscopic strain rate term in (2) plays a different role than the microscopic (i.e. junctional) strain rate feedback measured in [23].

### Mutant analysis validates machine-learned rules: Tissue flow requires an embryo-scale adhesion pattern

Our model (Box 2) predicts that tissue flow during GBE requires an embryo-scale gradient in cell adhesion (interpreted as the measured E-cadherin profile). We test this prediction using maternal mutants of *Drosophila* in which the DV patterning system is affected (Fig. 4). We compare the wild type (first row of the figure) to two mutants known as *spz*^4^ (second row) and *Toll*^*RM* 9^ (third row). These mutants adopt uniform cell fates corresponding to different positions along the DV axis of the wild-type embryo (arrows in first column of Fig. 4). Hence, the adhesion patterns are uniform along the DV axis (second column and panel b). Based on our machine-learned rules, we predict that no contractility gradient will form and no flow will occur in these mutants. Indeed, we observe that the myosin DV gradient is strongly suppressed in the mutants (third column and Fig. 4c), and no significant tissue flow is observed (fourth column and Fig. 4d). In both WT and mutant embryos, we observe an anti-correlation between E-cadherin and myosin levels in agreement with our learned rules. We note that in *Toll*^*RM* 9^ embryos, no flow occurs despite a uniformly high level of myosin, supporting the hypothesis that junctional myosin recruitment alone is not sufficient to drive tissue flow during GBE [21].

**Figure 4:**
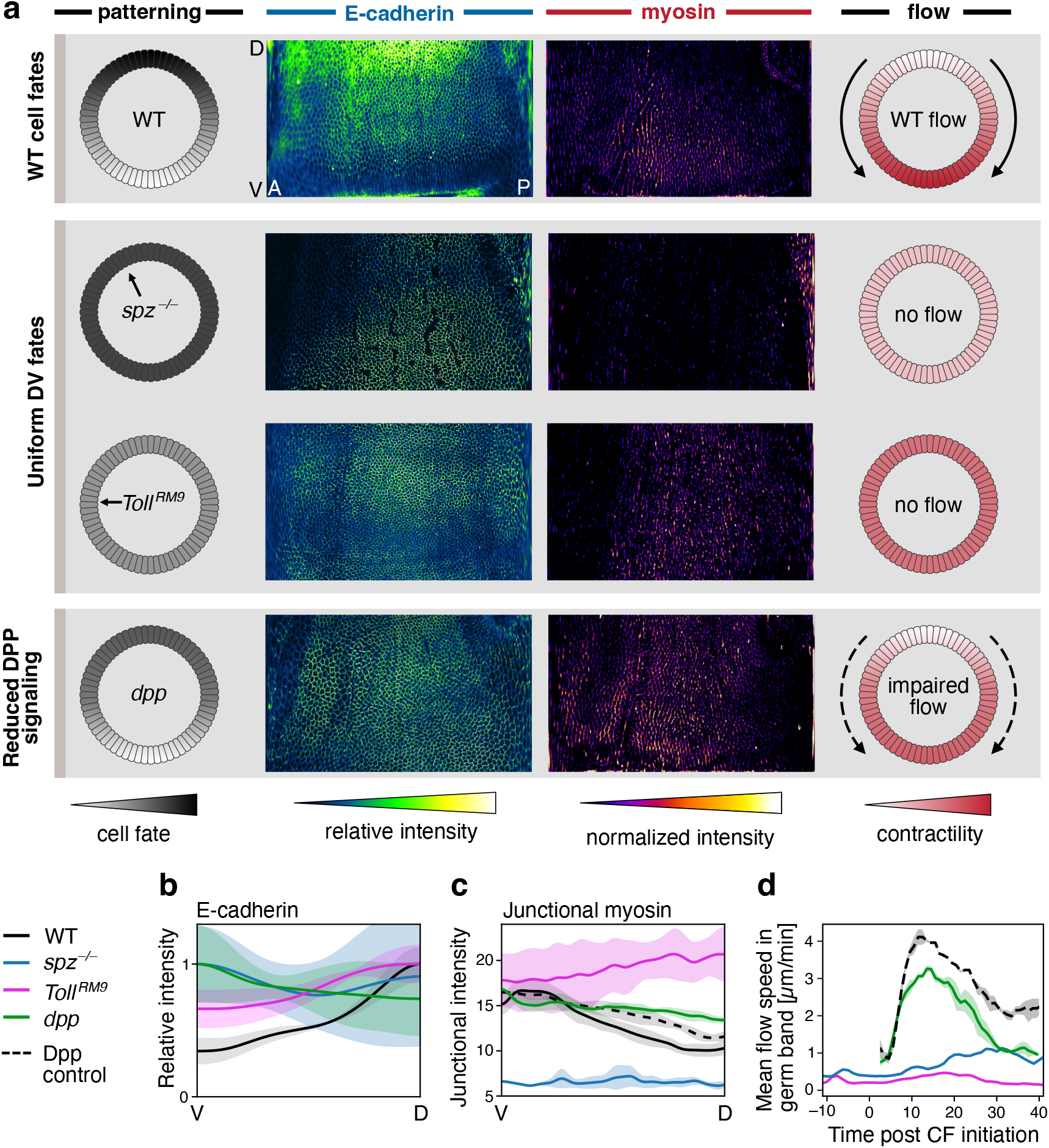
DV patterning mutants suppress cytoskeletal gradients and tissue flow. (**a**) (first column) DV-patterning “clock” cartoon of *Drosophila* gastrula. Color indicates differentiated cell fate. *Toll*^*RM* 9^ and *spz*^4^ embryos adopt uniform fates corresponding to positions marked with arrows. *dpp* mutants eliminate dorsal-most fates and lateral fates shift towards the dorsal pole. (second column) E-cadherin measured at initiation of the cephallic furrow. Fourth row shows E-cadherin in *dpp*^4^, *snail*^*IIG*05^ double mutants. (third column) Cytosolic-normalized myosin measured 15 min post initiation of the cephallic furrow. Fourth row shows myosin measured in *Dpp*^*hin*46*/*+^ mutants. (fourth column) Our model predicts that elimination of DV-patterned adhesion should impair or eliminate contractility gradients and tissue flow during GBE. (**b**) Quantification of E-cadherin gradient along DV axis for each genotype in (**a**). (**c**) Quantification of junctional myosin along DV axis for each genotype in (**a**). *Dpp*^*hin*46*/*+^ shows impaired grading. Data in (**b-c**) based on *N* = 3 embryos for each genotype. (**d**) Quantification of tissue flow in each genotype. Flow is suppressed in *Toll*^*RM* 9^ and *spz*^4^ mutants (imaged at 22° C). *Dpp*^*hin*46*/*+^ mutants impair but do not abolish tissue flow. *Dpp*^*hin*46*/*+^ and *Dpp*^+*/*+^ control embryos were imaged at 27° C which increases the speed of gastrulation over embryos imaged at lower temperatures [10].

### Zygotic regulation: BMP/Dpp signaling controls embryo-scale adhesion distribution

At this stage, our tentative picture is entirely post-translational. We now take a step back and ask: how does the embryo control the feedback loops involving only myosin and adherens junctions that we have identified? The embryo-scale DV gradient of E-cadherin is established before the onset of gastrulation, likely by a DV-graded morphogen. A natural candidate is Dpp, *Drosophila* homolog of BMP, a key morphogen responsible for differentiation of cell fate along the DV axis [38]. Dpp is graded like E-cadherin with highest concentration at the dorsal pole. Therefore, it could control the E-cadherin concentration early on. Reducing BMP signaling should then suppress the E-cadherin DV gradient, leading to a weaker contractility gradient and tissue flow. To test this, we analyzed *dpp*^*hin*46*/*+^ mutants (Fig. 4, fourth row), in which BMP signaling is reduced. We observe flattened E-cadherin and myosin patterns, as well as weaker flow relative to control embryos. This supports our hypothesis that BMP signaling establishes a DV-graded profile of E-cadherin and other components of the adherens junction.

### Signaling pathway linking AP to DV axes: BMP/Dpp role in patterning adhesion via PRGs

We now ask: what is the signaling pathway by which BMP establishes the embryo-scale adhesion pattern? Previous work has established that PRGs are involved in GBE (Box 1), often focusing on their prominent striped AP dependence [18–20]. Looking more closely, we observe that the expression levels of certain PRGs also exhibit gradients in the DV direction (Fig. 5b-d). Note also that Toll-6, a downstream target of the PRG Even-Skipped, does exhibit a strong DV dependence in WT embryos (Fig. 5e), even though the expression of Even-Skipped itself does not show a strong DV dependence (Fig. 5d). This motivates us to propose that BMP signaling may regulate E-cadherin through PRGs.

**Figure 5:**
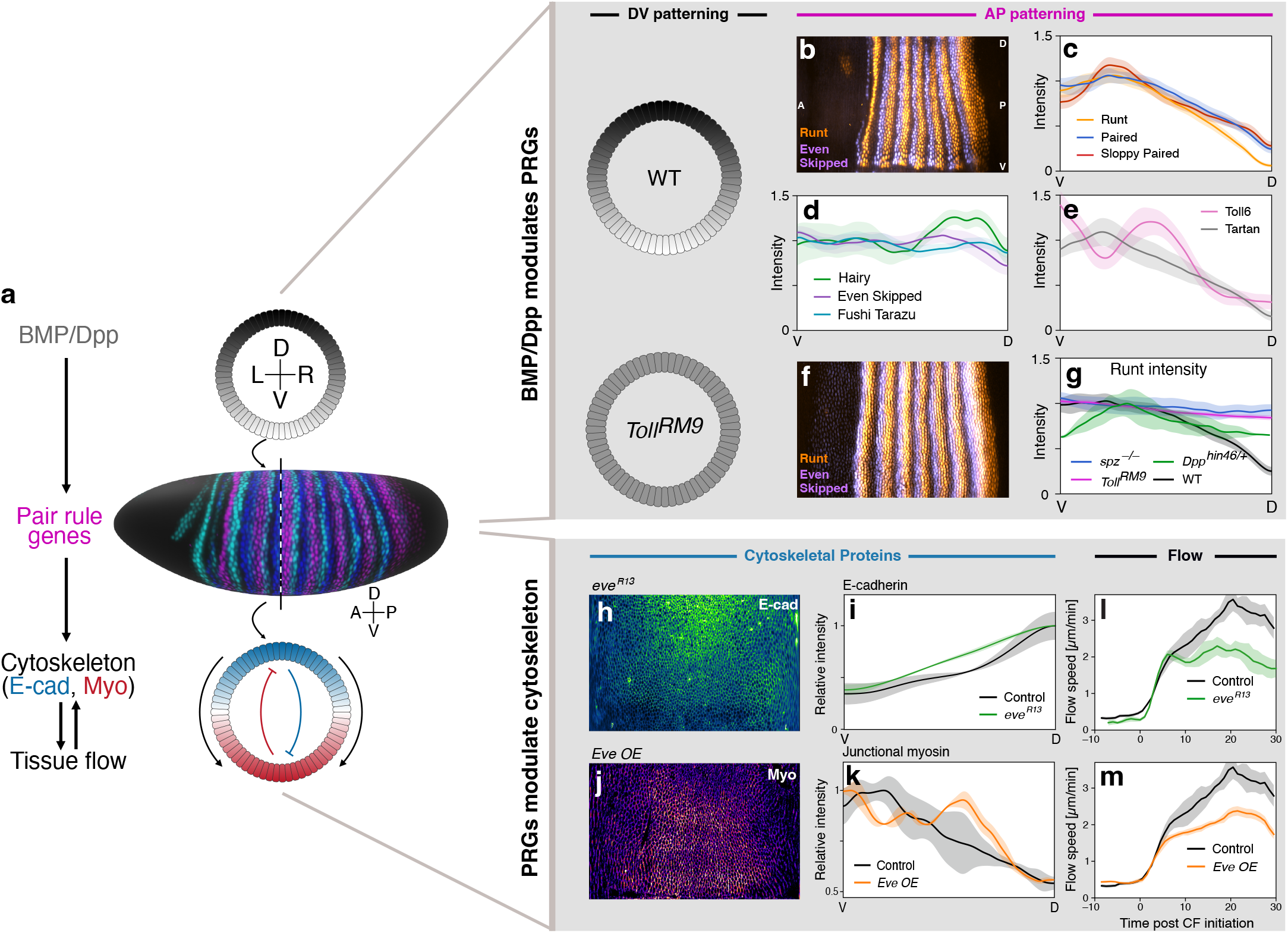
BMP signaling controls protein behavior via PRGs. (**a**) Cartoon of proposed signaling cascade. (**b-e**) In addition to stripes along the AP axis, the expression of certain PRGs exhibits a modulation along the DV axis. PRG expression along a central slice shows DV modulation for Runt, Paired, and Sloppy Paired (c) and weak modulation for Hairy, Even Skipped, and Fushi Tarazu (d). Downstream transmembrane receptors (Toll-6, Tartan) are also DV-graded (e). (**f**) *Toll*^*RM* 9^ mutants adopt uniform fates along the DV axis, but still exhibit AP stripes. Runt expression in these mutants shows no DV gradient. (**g**) Quantified Runt expression in DV mutants (Fig. 4) shows impaired or suppressed DV-modulation of myosin and E-cadherin. (**h-i**) *eve*^*R*13^ is a null allele of Even-skipped. E-cadherin expression in these mutants exhibits a shallower DV-gradient compared to control (*N* = 3 embryos). (**j-k**) *Eve OE* (*67,15*≫ *UAS-eve*) over-expresses Even-Skipped uniformly throughout the embryo. Myosin expression in *Eve OE* mutants is stronger near the dorsal pole of the embryo (*N* = 2 embryos). (**l-m**) Mutations to a PRG (*eve*^*R*13^ and *Eve OE* lead to weaker GBE flow (*N* = 3 embryos per genotype).

We test this hypothesis in two stages. To evaluate whether BMP/Dpp regulates PRG expression, we compare the PRG Runt in a WT embryo to the mutants *Toll*^*RM* 9^ and *spz*^4^ that we introduced in Fig. 4. While the AP-oriented stripes are unchanged, we observe that the DV gradient of Runt present in the WT disappears in the mutants (Fig. 5f-g).

To test whether PRGs regulate E-cadherin, we then turn to mutants where PRG expression is explicitly modified (see Methods). We observe that both inhibition of Even-Skipped via *eve*^*R*13^ mutants and upregulation via *67,15* ≫*UAS-eve* exhibit a weaker tissue flow compared to control embryos (Fig. 5l-m). In addition in *eve*^*R*13^ mutants, we observe reduction in the levels of E-cadherin in the middle of the DV axis (Fig. 5h-i). Consistently over-expression via *67,15*≫ *UAS-eve* shows an extension of the myosin pattern to the dorsal pole (Fig. 5j-k).

Together, these observations support the hypothesis that BMP signaling acts at least in part via the AP-patterned PRGs to establish a smooth DV-graded profile of E-cadherin expression before GBE. This gradient in cell adhesion then leads to a gradient in myosin recruitment along the DV axis, possibly by modulating mechanical feedback loops (see Box 2, Refs. [23, 39], and SI). The resulting contractility gradient drives tissue flow. A visual summary of the proposed mechanism is show in Fig. 5a.

### Conserved role of BMP and adhesion patterning from *Drosophila* to human neuroectoderm

BMP signaling has a conserved role in neuroectoderm fate determination from fly to vertebrates [7]. Molecular components of the cytoskeleton, such as myosin or E-cadherin, are also conserved between the species. Yet the morphogenesis of these tissues involves distinct shapes in different species. This raises the question of whether the post-translational mechanism we have identified in *Drosophila* (and summarized in Fig. 5a) can be conserved in organisms with different geometries.

To address this question, we turn to a reproducible human stem cell-based model of neural tube morphogenesis. In this model, stem cell cultures with a single lumen, controlled geometry, volume, and cell number are differentiated into neuroectoderm by neural induction and timed BMP4 supplement (Fig 6a), see Methods and Refs. [40, 41]. In contrast with *Drosophila*, where the germband extends along the surface of an ellipsoid but does not fold, in humans the neural tube folds on the surface of a disc. BMP signalling is driven from the edge, and the resulting gradient organizes patterned differentiation into surface ectoderm (in blue in Fig 6b) and neural ectoderm (in purple in Fig 6b) [42]. In this stem cell-based model, the neural tube undergoes major shape changes, including bending, folding and closure [40]. These major shape changes have important consequences on mechanics, and may complicate the interpretations of cytoskeletal patterns. Hence, we focus on the early configuration of the cytoskeleton, that triggers the flows leading to bending (i.e. before 36 hours post BMP supplementation (hpBMP), see Methods).

**Figure 6:**
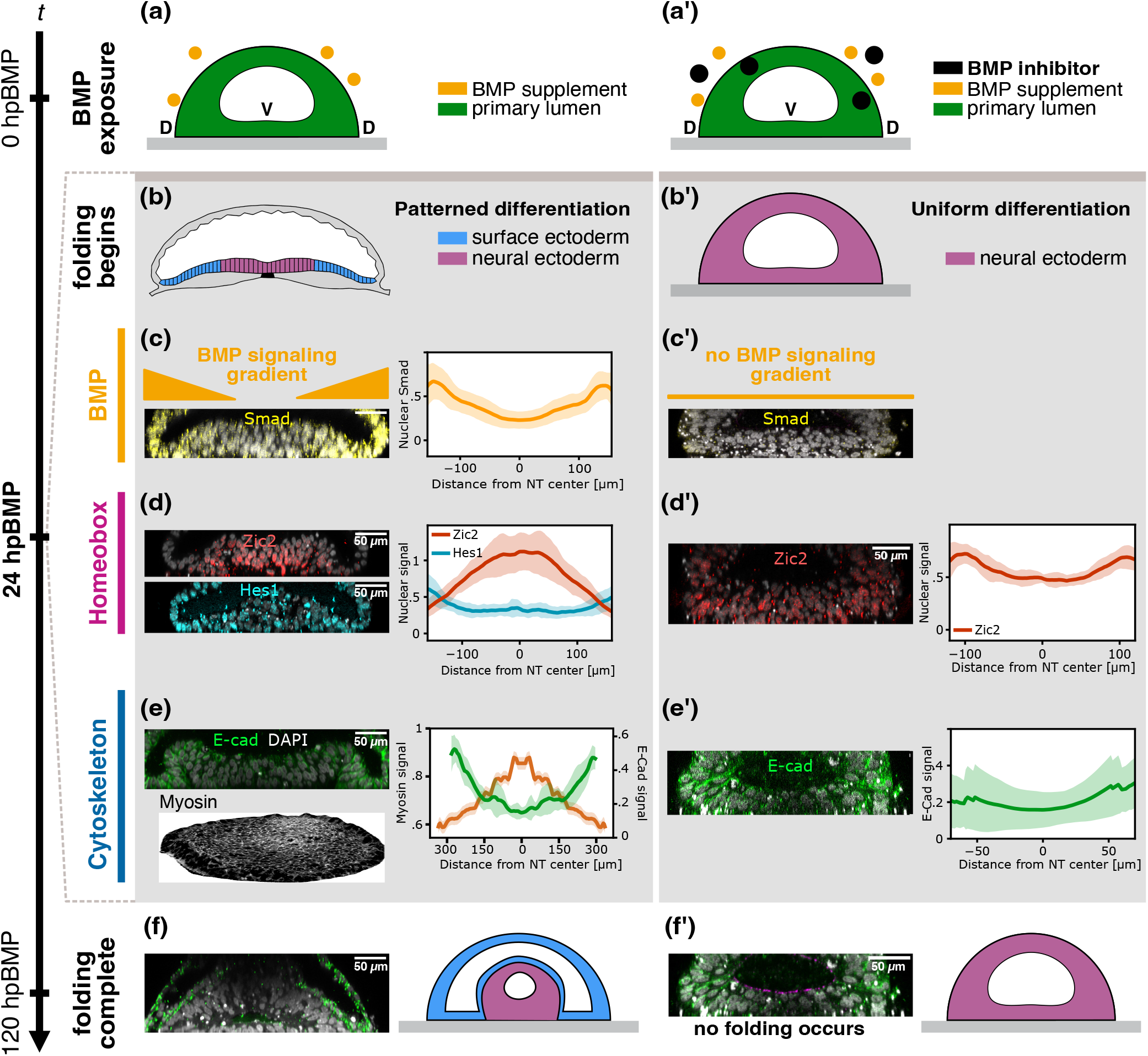
Patterned expression in human neural tube organoids (NTO). (**a**) 3D stem cell sheets are exposed to BMP4 to drive tissue folding. All times are measured as hours post-BMP exposure (hpBMP). (**b**) By *t* = 24 hpBMP, the tissue differentiates into surface and neural ectoderm and first morphogenetic movements begin. (**c**) Left: Nuclear SMAD 1/5/9 (Smad), a BMP4 signaling marker, is used to readout BMP4 signaling. NTO stained for Smad shows an expression gradient, taken at *t* = 20 hpBMP. Right: Quantification obtained from the average nuclear signal across *z*-slices in NTO (surface ectoderm dome excluded) in *N* = 6 NTOs). (**d**) Left: Zic2 and Hes1 are homologs of *Drosophila* PRGs. NTOs stained for Zic2 and Hes1 exhibit expression gradients in relation to the SMAD gradient, taken at *t* = 20 hpBMP. Right: Quantification obtained using *N* = 6 NTOs. (**e**) Left: E-cadherin, taken at *t* = 20 hpBMP, and apical myosin, taken at *t* = 24 hpBMP, expression in NTOs. Both exhibit a gradient in relation to Smad. Right: Quantification obtained using *N* = 6 NTOs for E-cad. and *N* = 2 NTOs for myosin. (**f**) The folding process is complete by *t* = 120 hpBMP. (**a’-f’**) Repeated analysis on NTOs treated with BMP4 inhibitor LDN (200*n*M). The resulting NTO lacks surface ectodermal differentiation (**b’**) as signified by absense of Smad straining (**c’**), homebox and E-cadherin expression profiles exhibit no significant gradients (**d’-e’**), and no folding occurs (**f’**).

We find that all the elements present in our proposed mechanism for *Drosophila* (Fig. 5a) also show organ-scale gradients in the stem cell-based model, and mirror BMP signaling. By 20 hpBMP we observe a stable BMP signaling gradient (Fig. 6c). At the same time, we see differential expression patterns of homeobox genes such as Zic2 and Hes1, homologs of *Drosophila* PRGs (Fig. 6d). Adherens junction proteins such as E-cadherin are also smoothly graded at this time, and only become sharply delineated from the neural ectoderm later. By 24 hpBMP, myosin exhibits a smooth gradient that counter runs the E-cadherin profile (Fig. 6e). We note that because the geometry of the human *in vitro* neural tube differs from that of *Drosophila*, there is no geometric stress (Box 2) to drive local anisotropic recruitment of myosin. Nevertheless, the comparison with *Drosophila* suggests that both force-generating proteins and PRG orthologs are modulated by the BMP signaling gradient, suggesting that this role of BMP in setting the initial condition for the morphogenetic events is conserved.

To test this further, we eliminate BMP signaling via the BMP inhibitor LDN (Fig 6a’), so that there is no BMP gradient (Fig 6c’). As expected, all cells differentiate into neural ectoderm: the resulting structures completely lack surface ectoderm (compare Fig 6b and b’). We further observe that, as a consequence, PRG orthologs and E-cadherin also exhibit no gradient (Fig. 6d’,e’). Finally, the structures where BMP signaling is suppressed do not fold (Fig 6f’). This supports our hypothesis that the organoid-scale gradient in BMP signaling primes the cytoskeleton to orchestrate the morphological movements that initiate neural tube closure, possibly through PRG homologs.

To sum up, despite striking differences in shape, the *in vitro* neural tube model and *Drosophila* share a crucial geometrical feature: there is a key axis defined by BMP patterning in both systems, along which a contractility gradient gets established. Our results suggest that this general feature is conserved, but fine-tuned by the geometry of different organisms.

## DISCUSSION

In this work, we have proposed a global picture of neuroectoderm morphogenesis which links the conserved mechanisms of BMP signaling and convergent extension. Figure 7 summarizes how this picture materializes in *Drosophila* neuroectoderm morphogenesis (left) and in a human neural tube model (right). In both cases, BMP signaling gradients are relayed into a global pattern in adherens junction proteins via genetic regulation. This in turn sets up a global pattern of force-generating cytoskeletal proteins, such as myosin, that modulate tissue mechanics. In *Drosophila*, this arises through mechanical feedback loops involving tissue-scale stress and external strains, which result in coordinated morphogenetic flow [22, 23]. Whether human neural tube closure relies on similar feedback mechanisms for governing protein dynamics remains an open question, but we note that Zic family proteins are involved in neural tube morphogenesis [43]. This overall bio-mechanical picture might be conserved even beyond bilaterians. In *Nematostella* (sea anemones), for example, BMP signaling regulates a distinct set of homeobox genes [44] also associated with the mechanical processes sculpting the body [45].

**Figure 7:**
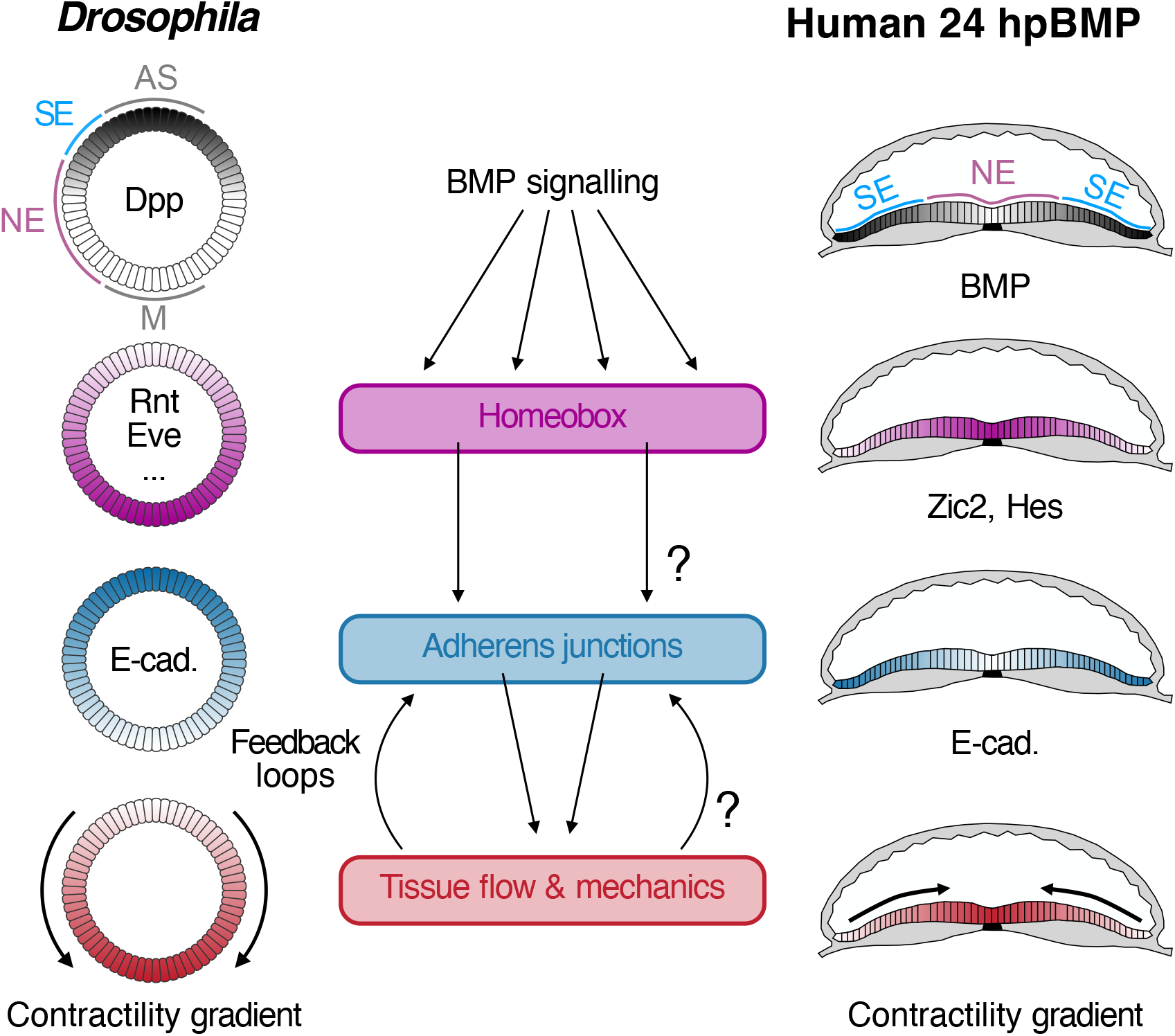
Proposed conserved signaling cascade from BMP signaling to tissue flow. Hypothesized morphogenetic parallels between *Drosophila* gastrulation and human neural tubes. SE: surface ectoderm, NE: neural ectoderm, M: mesoderm, AS: amnioserosa. DPP/BMP signaling establishes a gradient in Homeobox transcription factors (PRGs in *Drosophila*), which lead to large scale cytoskeletal gradients, notably of E-cadherin and adherens-junctional proteins. These gradients lead to tissue flow, which can feed back to the cytoskeleton. Open questions in the neural tube organoids are the causal role of homeobox gradients, and the presence or absence of mechanical feedback loops.

The picture we have developed aligns with classical work showing that inverting the BMP gradient via RNA injection reverses the direction of *Drosophila* neuroectoderm extension [46]. However, it is complementary to the heterotypic juxtaposition model [9]. Instead of direct genetic regulation locally modulating cell adhesion and anisotropic forces at cell boundaries, our model emphasizes the role of a embryo-scale modulation of adherens junctions which breaks the DV symmetry in a manner prescribed by BMP signalling. A molecular mechanism for this large-scale modulation of adherens junction proteins may involve Toll-like transmembrane receptors (TLRs; see Fig. 5b). The precise mechanism by which TLRs regulate the cytoskeleton remains incompletely understood [9]. However, TLRs are known to strongly impact neural epithelium elongation in *Drosophila* and to be controlled by PRGs [47, 48].

The picture of neuroectoderm morphogenesis proposed in this work has been obtained by developing a set of causal rules, that we then tested in experiments. These causal rules were identified by complementing data-driven inference with insights from biology and physics. This is not a new way of doing biology. Nonetheless, our use of machine learning techniques aids the formulation and testing of biological hypothesis and mathematical models [17]. What we can learn also depends on the interpretative framework that we use. For instance, our coarse-grained picture neglects the details of cell-level mechanisms. As a result, our model shows discrepancies with experimental measurements of mechanical feedback at strained junctions (Box 2 and SI), which are relevant during ventral furrow invagination [23].

We conclude by asking: why should the mechanisms by which morphogenesis unfolds be conserved? In biology, usual sources of universality are the conservation of genes by natural selection and convergent evolution. Morphogenesis has the peculiarity of strongly mixing biological and mechanical constraints: organisms have to abide to the laws of mechanics to get their shape. We conjecture that these physical constraints translate into evolutionary constraints that make morphogenesis more likely to exhibit conserved mechanisms. Our approach may accelerate the systematic identification of such universal mechanisms – conserved across species – that biology uses to seed the shape of organisms.

## Acknowledgments

We thank Susan Wopat, Aimal Khankhel, and the members of the Streichan lab for helpful discussions. This work was supported by grants NIGMS R35-GM138203 and CAREER PHY:2047140. JC and MF acknowledge support from the National Science Foundation under grant DMR-2118415. MF and VV acknowledge partial support from the UChicago Materials Research Science and Engineering Center (NSF DMR-2011864). VV acknowledges support from the Army Research Office under grant W911NF-22-2-0109 and W911NF-23-1-0212 and the Theory in Biology program of the Chan Zuckerberg Initiative. This research was supported from the National Science Foundation through the Center for Living Systems (grant no. 2317138). This work was completed in part with resources provided by the University of Chicago’s Research Computing Center.

## Notes

### Competing Interest Statement

The authors have declared no competing interest.

